# Sampling design and inference of the caecal–skin *Campylobacter* relationship in broilers

**DOI:** 10.64898/2026.05.03.722495

**Authors:** Celine Mason, Erica Nunney, Javier Guitian

## Abstract

The relationship between Campylobacter levels in broiler caeca and on carcass skin is central to quantitative microbial risk assessment along the poultry production chain, underpinning modelling of intervention impacts, including EFSA assessments of the public health impact of control measures. However, this relationship is typically inferred from monitoring data generated under sampling designs that do not preserve pairing between specimens and may involve pooling. In this study, we used a simulation framework to evaluate whether commonly used sampling strategies allow reliable recovery of the caecal-skin relationship.

A simulated broiler population was generated, assigning caecal and skin loads to individual birds based on a specified linear relationship. Sampling was conducted under paired and unpaired designs, with and without pooling, reflecting approaches used in surveillance programmes and in policy-oriented models. Regression models were fitted to sampled data across 1,000 simulations for a range of assumed slopes.

Under paired sampling, estimated slopes closely matched the true relationship across most scenarios. In contrast, unpaired sampling consistently failed to recover the association, with estimated slopes centred around zero regardless of the true slope. These findings were robust to variation in within-flock prevalence, residual error, and intercept.

The results show that sampling design fundamentally affects identifiability of relationships between stages of the production chain. This has implications for interpretation of parameters derived from monitoring data and used in quantitative Campylobacter risk assessments informing policy. Parameters derived from unpaired and pooled monitoring data should therefore be interpreted with caution when used to support risk assessment and decision-making.

Campylobacter; broiler chickens; sampling strategy; unpaired sampling; carcass contamination; quantitative microbial risk assessment; simulation.

## Introduction

*Campylobacter* spp. are the most common bacterial cause of foodborne gastrointestinal disease in the EU and the UK (EFSA BIOHAZ Panel, 2020), with poultry recognised as a major reservoir and source of human infection (Kintz, 2024). Campylobacter infection is typically associated with self-limiting gastrointestinal illness but can lead to severe complications such as Guillain–Barré syndrome (Finsterer, 2022). The main species of concern for human health are *Campylobacter jejuni* and *Campylobacter coli* (ECDC, 2024).

In 2010 the UK Joint Working Group on Campylobacter set a target to reduce the proportion of highly contaminated post-chill chicken carcasses (> 1,000 CFU/g) from 27% to 10% by 2015 based on an European Food Safety Authority (EFSA BIOHAZ Panel, 2010) scientific opinion suggesting that a 10% reduction in carcass contamination would reduce public health risk by 50-90% (EFSA BIOHAZ Panel, 2011; Wearne, 2015). Despite a substantial reduction in highly contaminated chicken at retail, the expected decrease in human cases was not observed, remaining stable at around 100 cases per 100,000 population between 2014-2019 (UKHSA, 2025). More recently, reported campylobacteriosis cases have increased, with a 17.1% rise in England in 2024 compared with the previous year (UKHSA, 2025), consistent with observations that reductions in poultry contamination have not necessarily translated into declines in human cases in other settings (Kintz et al. 2024).

In broilers, Campylobacter primarily colonises the caeca and small intestine without causing clinical signs and carcass contamination occurs during slaughter through faecal leakage and cross-contamination (Newell & Fearnley, 2003; Josefsen, 2015). Measuring the levels of Campylobacter contamination in broilers involves sampling different specimen types taken at different points of processing, including caecal contents at evisceration and neck skin post chill (Dubovitskaya et al, 2023). Monitoring programs often collect multiple caecal and carcass samples per slaughter batch without bird-level matching between specimen types. For example, the UK monitoring programme (2016–2017) collected ten caecal samples at the point of evisceration alongside one post-chill carcase sample (neck and breast skin) (Lawes, 2019).

In 2020 EFSA published an updated assessment of control options for Campylobacter in broilers, including quantitative modelling of the expected impact of different control options on reducing Campylobacter spp. contamination in carcases (EFSA BIOHAZ Panel, 2020). As in the 2011 opinion, EFSA used a linear regression approach to relate Campylobacter concentrations in chicken caeca to concentrations on broiler skin, used as a proxy for contamination on meat after processing. This approach was adopted as a pragmatic alternative to explicitly modelling the complex dynamics of Campylobacter transfer and survival during industrial processing. The uncertainty in this relationship was represented through a Beta-pert distribution for the slope parameter, with a most likely value of 0.27, a minimum of 0, and a maximum of 0.7. The most likely value derived from the largest dataset from UK monitoring data generated under pooled and unpaired sampling, in which ten pooled caecal samples and one neck/breast skin sample were collected per batch (Lawes, 2019). An implicit assumption in modelling approaches such as that adopted by EFSA, which link Campylobacter concentrations in the caeca to those measured on carcass skin, is that the underlying association is identifiable from the available data. However, monitoring sampling schemes are typically designed to estimate contamination levels rather than to preserve bird-level or batch-level pairing between caecal and skin samples. Under such designs, the structure of the sampled data itself may limit the ability to infer the true relationship between caecal and skin contamination. This is consistent with the heterogeneity reported in the published literature available to EFSA at that time: of 15 studies identified, six reported no significant correlation between caecal and skin or meat concentrations, while the remaining studies reported slope values ranging from 0.21 to 1.15. Even among the larger studies, findings were inconsistent, with one reporting no positive correlation and others reporting slopes between 0.21 and 0.32.

In particular, unpaired and pooled sampling strategies may fundamentally alter the information required to identify this relationship. This raises a fundamental question of identifiability: whether the underlying relationship between caecal and skin contamination can be inferred from the available data.

We hypothesise that, under commonly used sampling strategies, the caeca–skin relationship may not be reliably recoverable from the observed data. Using a simulation framework, we evaluate how different sampling designs influence bias and identifiability in estimation of the caecal–skin relationship, with particular reference to sampling schemes used to derive parameters for quantitative microbial risk assessment models informing policy.

## Methods

### Overview of the modelling approach

A simulated broiler population was created in which each chicken was assigned a level of caecal and skin Campylobacter load. Caecal loads were generated by assuming a fixed within-flock prevalence and a normal distribution of log10-transformed bacterial loads among colonised birds. Skin loads were then inferred from caecal loads using a linear regression of log10-transformed neck skin on caeca values.

For each scenario, the simulated population was sampled under alternative designs reflecting monitoring approaches used in routine surveillance, including paired and unpaired sampling, with and without pooling. In pooled designs, a single composite caecal sample was formed from multiple specimens, whereas in unpaired designs caecal and skin samples were taken from different birds within the same flock. A linear regression was then fitted to the sampled data to estimate the observed slope relating caecal and skin contamination, analogous to the approach used in the EFSA quantitative modelling of intervention impact (EFSA BIOHAZ Panel, 2020).

The sampling and estimation procedure was repeated 1,000 times to generate a distribution of sampled regression slopes. This allowed evaluation of whether the underlying caecal-skin relationship specified in the simulated population could be recovered under different sampling designs.

### Simulation of the broiler population

The population consisted of 100 flocks of 20,000 birds each. Each bird was assigned a colonization status (colonized or not colonized) using a binomial distribution, with within flock prevalence (wfp) as the probability of a bird being colonized. The wfp was fixed at 70% (Bull et al., 2008) as a baseline value; its influence was explored in subsequent scenario analyses. Uncolonized birds were assigned 1 CFU/g (log10 = 0) to avoid undefined values following log transformation. Infected birds were given a caecal Campylobacter concentration in log10 CFU/g that is randomly drawn from a normal distribution with the specified caeca mean (5.94) and standard deviation (3.71). These distribution parameters were derived from the APHA/FSA monitoring programme data (Rodgers, 2020).

For each bird, skin Campylobacter concentration was then generated from caecal concentration using the linear relationship:

log_10_ *skin* = *a log*_10_ *caeca* + b + *ε*

where:

*a* is the slope of regression line (true slope),

*b* the intercept and

*ε* is a normally distributed error term with mean of 0 and standard deviation σ.

In the model, the residual standard deviation (σ) was set to 0.1 log10 CFU/g, and was varied in scenario analyses to assess the impact of residual variability.

The simulation was carried out for different assumed values of the slope of the regression line (true slope), therefore generating a distribution of sampled slopes (observed slopes) conditional on the assumed true slope. Parameters used to define the simulated population are shown in Table 1.

**Table 1.** Parameters defining the simulated population and the caecal colonization–carcass contamination model.

| Parameter | Value | Description | Source |
| --- | --- | --- | --- |
| Caecal mean | 5.94 log<sub>10</sub> CFU | Mean of log<sub>10</sub>-transformed caecal load distribution in colonised birds | Rodgers, 2020 |
| Caecal standard deviation | 3.71 | Standard deviation of log<sub>10</sub>-transformed caecal load distribution | Rodgers, 2020 |
| Skin intercept | 0 | Intercept of the assumed linear relationship between caecal and skin contamination | Assumed |
| Error (residual SD) | 0.1 | Standard deviation of residual error term in caecal-skin relationship (log<sub>10</sub> scale) | Assumed |
| Detection limit | 5 CFU/g | Lower detection limit; values below threshold assigned random values between 1 and detection limit | Lawes, 2019 |

Caecal loads were generated on the log10 scale and transformed to CFU/g for pooling and application of the detection limit. Following these steps, values were log10-transformed prior to regression analysis.

### Sampling strategy

Four different sampling scenarios were explored: Paired Individual; Paired Pooled; Unpaired Individual; Unpaired Pooled. Each scenario was sampled independently from the simulated population.

In paired scenarios, caecal and skin bacterial loads were obtained from the same birds, thereby preserving matched observations.

In unpaired scenarios, caecal and skin loads were sampled independently from birds within the same flock.

The UK monitoring programme was used as the basis for an unpaired sampling scenario, in which one carcass (neck and breast skin) sample and one composite sample of 10 caeca per slaughter batch were collected (Smith et al., 2023).

Paired scenarios were fixed at ten caeca and ten skin samples per flock, and pooled scenarios were set at ten caeca and one skin sample per flock.

In pooled scenarios, concentrations were averaged on the CFU/g scale to represent composite samples, whereas individual scenarios retained bird-level measurements.

To reflect routine microbiological testing, a detection limit of 5 CFU/g was applied after pooling, consistent with the UK monitoring programme (FS101126; Lawes, 2019). Samples below 5 CFU/g were replaced by a value drawn from a uniform distribution between 1 and 5 CFU/g to represent measurement uncertainty below the detection limit.

### Caecal-skin regression model

For each simulated dataset, a simple linear regression was fitted on the log10 CFU/g scale to quantify the observed relationship between caecal and skin Campylobacter concentrations under the given sampling strategy and assumed true regression slope. The estimated regression coefficient was referred to as the sampled slope.

The simulation was repeated 1,000 times for each specified true slope.

### Statistical analysis

For each sampling scenario, the median and mean sampled slope over all simulations was recorded. A reference slope of 0.276 was used as a benchmark, as this value, derived from the APHA/FSA monitoring programme for Campylobacter in broilers (2012-2017) (Rodgers, 2020), formed the basis of the most likely slope value of 0.27 adopted in the EFSA modelling approach (EFSA BIOHAZ Panel, 2020). The proportion of simulations in which the estimated slope fell within ±0.05 of the true slope and within ±0.05 of the target slope (0.276) was obtained.

### Scenario analysis

The simulation was repeated to evaluate the robustness of results to key assumptions. First, within flock prevalence was varied across four levels: 30%, 50%, 70% (baseline) and 100%. Second, the residual error term used to generate skin contamination from caecal loads was varied to assess the impact of measurement variability. In addition to the baseline value of 0.1 log10 CFU/g, simulations were repeated with standard deviations of 0.5 and 1.0 log 10 CFU/g. Third, sensitivity to the intercept parameter of the caecal-skin relationship was explored by repeating simulations with intercept values of 0, 1, 10, -1 and -10, while keeping within-flock prevalence fixed at 70%. For each scenario, the same sampling and estimation procedures were applied, and summary statistics of the sampled slopes were obtained as described above.

### Reproducibility

Analyses were conducted in R (version 4.4.2) using the packages dplyr, ggplot2, tidyr, magrittr, and roxygen2. Simulations were performed with 1,000 iterations for each scenario. A fixed random seed (42) was used to ensure reproducibility. All simulation code and outputs used in this study are available from the authors upon request.

## Results

The distribution of sampled slopes and the proportion of simulations in which the true and target slopes were recovered are shown in Table 2 and Figure 1. Under paired sampling strategies, the mean and median sampled slopes closely matched the true slope across most scenarios (Figure 2). In contrast, under unpaired strategies both the mean and median sampled slopes were approximately zero for all true slopes.

**Figure 1.**
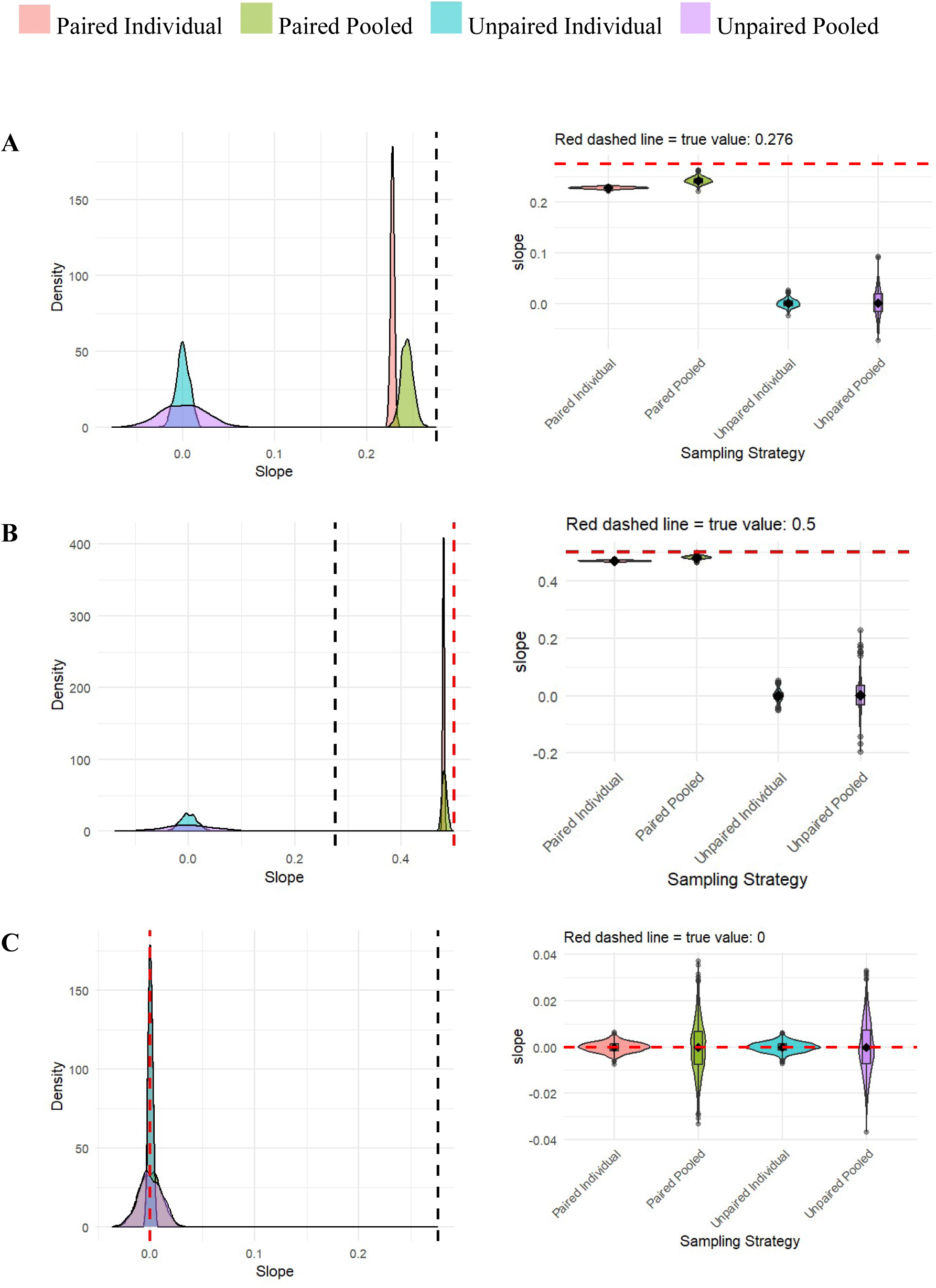
Distribution of sampled slopes under different sampling strategies. Density plots (left) and summary distributions (right) are shown for true slopes of: A = 0.276; B = 0.5; C = 0. Red dashed line indicates the true slope; grey dashed line indicates the target slope (0.276).

**Figure 2.**
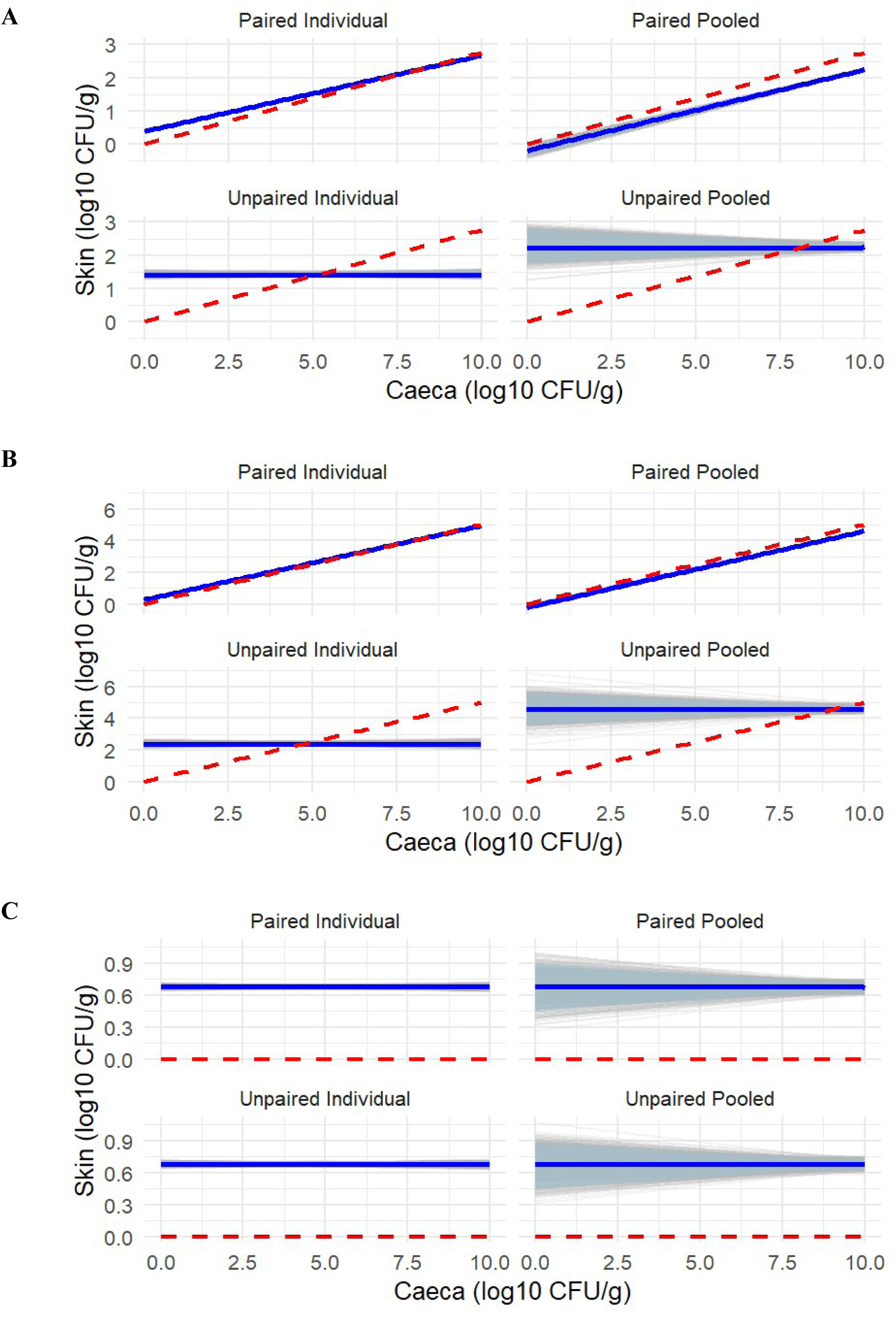
Estimated caecal–skin relationships under different sampling strategies. Panels show regression lines fitted to sampled data for true slopes of: A = 0.276; B = 0.5; C = 0. Grey lines represent individual simulations, the blue line the mean estimate, and the shaded area the 95% confidence interval. The red dashed line indicates the true slope. Note that y-axis scales differ between panels to aid visualisation of variation across scenarios.

**Table 2.** Results of sampled slopes for 1,000 simulations for different true slopes (−1, -0.5, 0, 0.276, 0.5, 1, 10) and four different sampling strategies (Paired individual, Paired pooled, Unpaired individual, Unpaired pooled). The mean and median of the sampled slopes is shown for each strategy. The percentage of scenarios with a sampled slope within a tolerance of 0.05 of the true slope and the target slope (0.276) are shown.

| True slope | Sampling strategy | Mean sampled slope | Median sampled slope | Sampled slopes within 0.05 of true slope (%) | Sampled slopes within 0.05 of target slope of 0.276 (%) |
| --- | --- | --- | --- | --- | --- |
| 0 | Paired individual | 0.000 | 0.000 | 100 | 0.00 |
|  | Paired pooled | 0.000 | -0.001 | 100 | 0.00 |
|  | Unpaired individual | 0.000 | 0.000 | 100 | 0.00 |
|  | Unpaired pooled | 0.000 | 0.000 | 100 | 0.00 |
| 0.276 | Paired individual | 0.228 | 0.228 | 86.7 | 86.7 |
|  | Paired pooled | 0.243 | 0.243 | 99.9 | 99.9 |
|  | Unpaired individual | 0.000 | 0.000 | 0.00 | 0.00 |
|  | Unpaired pooled | 0.001 | 0.001 | 0.00 | 0.00 |
| 0.5 | Paired individual | 0.468 | 0.468 | 100 | 0.00 |
|  | Paired pooled | 0.481 | 0.481 | 100 | 0.00 |
|  | Unpaired individual | 0.000 | 0.000 | 0.00 | 0.00 |
|  | Unpaired pooled | 0.001 | 0.003 | 0.00 | 0.100 |
| 1 | Paired individual | 1.00 | 1.00 | 100 | 0.00 |
|  | Paired pooled | 1.00 | 1.00 | 100 | 0.00 |
|  | Unpaired individual | 0.000 | -0.001 | 0.00 | 0.00 |
|  | Unpaired pooled | 0.000 | 0.001 | 0.00 | 1.70 |
| 10 | Paired individual | 10.52 | 10.52 | 0.00 | 0.00 |
|  | Paired pooled | 10.14 | 10.14 | 0.30 | 0.00 |
|  | Unpaired individual | -0.007 | -0.015 | 0.00 | 9.10 |
|  | Unpaired pooled | 0.003 | 0.013 | 0.00 | 5.50 |

This pattern arises because, under unpaired sampling, caecal and skin measurements are drawn from different birds, thereby removing the covariance structure linking the two variables. As a result, even when a strong relationship exists at the individual level, the sampled data do not retain this association, and regression estimates converge towards zero. This holds even when all birds are colonised and residual variability is low, as unpaired sampling breaks the correspondence between paired measurements rather than altering their marginal distributions.

For paired strategies, more than 85% of sampled slopes fell within 0.05 of the true slope (except for the scenario with a true slope of 10), indicating consistent recovery of the underlying relationship. Under unpaired strategies, no sampled slopes fell within this range for any non-zero true slope. Under paired sampling, pooled designs yielded estimates that were at least as close to the true slope as individual sampling, reflecting reduced variability in averaged measurements.

In the scenario analysis, varying within-flock prevalence had limited impact on slope estimation under paired sampling, although lower prevalence reduced precision. Under unpaired sampling, sampled slopes remained centred around zero across all prevalence levels.

Increasing the residual error reduced precision under paired sampling, with a lower proportion of estimates close to the true slope, while results under unpaired sampling remained centred around zero.

Variation in the intercept had minimal impact on slope estimation under paired sampling. Under unpaired strategies, sampled slopes remained centred around zero regardless of intercept value. Full results of the scenario analysis are provided in Supplementary Table S1 and Figure S1.

## Discussion

The relationship between Campylobacter concentrations in broiler caeca and on carcass skin is central to quantitative assessments of foodborne risk and has been used to inform predictions of the public health impact of farm-level control measures. However, this relationship is typically inferred from monitoring data generated under sampling designs that do not preserve pairing between specimen types and may involve pooling. In this study, we used a simulation framework to evaluate whether such sampling strategies allow reliable recovery of the underlying caeca-skin relationship. Our results show that, while paired sampling enables accurate estimation of the true relationship, unpaired designs consistently fail to recover it, with estimated slopes centred around zero under unpaired sampling, regardless of the true underlying association.

These results highlight that identifiability of the caeca-skin relationship depends critically on the sampling design. When measurements are linked at the bird level, the underlying association can be reliably inferred. In contrast, under unpaired sampling, the relationship is not recoverable from the observed data, irrespective of the true underlying slope. This indicates that the loss of identifiability is a structural consequence of the sampling design rather than a result of specific parameter choices or biological variability.

These findings have direct implications for the interpretation of parameter values used in EFSA modelling of the public health impact of interventions (EFSA BIOHAZ Panel, 2020). The slope value of 0.276, derived from UK monitoring data generated under pooled and unpaired sampling (Lawes, 2019; EFSA BIOHAZ Panel, 2020), is unlikely to reliably represent the biological relationship between caecal and skin contamination. Rather, the results suggest that such estimates may arise from sampling designs that do not preserve the underlying association, leading to biased and potentially misleading estimates of the relationship. As a result, predictions of the public health impact of interventions based on this relationship should be interpreted with caution, as they may reflect artefacts of the data structure rather than the true effect of changes in caecal colonisation on carcass contamination (EFSA BIOHAZ Panel, 2020). This has implications for the interpretation of model-based estimates of the public health impact of on-farm interventions, which rely on the caecal-skin relationship. More broadly, it highlights the importance of accurately characterising relationships between stages of the production chain when linking changes in contamination levels to human health outcomes, in light of the discrepancies between trends in contamination and human campylobacteriosis reported in recent evaluations (Kintz et al., 2024).

From a practical perspective, these results highlight important limitations of commonly used monitoring approaches for informing quantitative risk assessments. Sampling strategies based on unpaired, and often pooled, specimens may be suitable for estimating contamination levels but may not be reliable for inferring relationships between stages of the production chain.

Where such relationships are required to support modelling or evaluation of interventions, sampling designs that preserve pairing between specimen types should be considered, in study designs specifically aimed at estimating relationships between contamination stages.

In routine surveillance contexts where this is not feasible, the use of such data for estimating inter-stage relationships should be approached with caution.

The variability in findings reported in the literature regarding the relationship between caecal and carcass contamination may be explained, in part, by differences in sampling design.

Some studies have reported strong associations between caecal and carcass contamination, while others have found weak or no relationship (Allen et al., 2007; Boysen et al., 2016; Reich et al., 2008). The results of the present study suggest that such discrepancies may arise from differences in the extent to which sampling strategies preserve pairing, rather than reflecting true differences in the underlying biological relationship.

From a methodological perspective, these findings point to the need for study designs that preserve pairing between caecal and skin samples at the bird level. An ideal approach would involve collecting matched caecal and carcass samples from the same birds across a range of contamination levels and production settings, allowing direct estimation of the relationship while accounting for biological and processing variability. While such designs may not be feasible in routine surveillance, they could be implemented in targeted studies to support parameterisation of risk assessment models.

This study has several limitations. The analysis is based on a simulation framework and therefore depends on assumptions regarding the distribution of contamination levels and the form of the relationship between caecal and skin contamination. In addition, a relatively simple linear relationship was assumed, and other sources of variability present in real production systems were not explicitly modelled. However, the consistency of the results across a wide range of parameter values supports the robustness of the main conclusions regarding the impact of sampling design on identifiability.

In conclusion, this study shows that commonly used sampling designs may not preserve the information required to infer relationships between stages of the production chain. Using Campylobacter in broilers as an example, and in the context of persistently high human infection rates, unpaired and pooled sampling can produce estimates that do not reflect the underlying association. These findings highlight the need to align sampling design with analytical objectives in quantitative microbial risk assessment.

## CRediT authorship contribution statement

**Celine Mason:** Writing – original draft, Writing – review and editing, Formal analysis, Methodology, Data curation.

**Erica Nunney:** Methodology, Writing – review and editing.

**Javier Guitian:** Conceptualization, Methodology, Writing – review and editing, Supervision.

## Declaration of generative AI and AI-assisted technologies in the manuscript preparation process

During the preparation of this work, the authors used ChatGPT (OpenAI) to assist with language editing and refinement of the manuscript text, and to support code review. The authors reviewed and edited the content as needed and take full responsibility for the content of the published article.

## Supplementary Material

**Figure S1.**
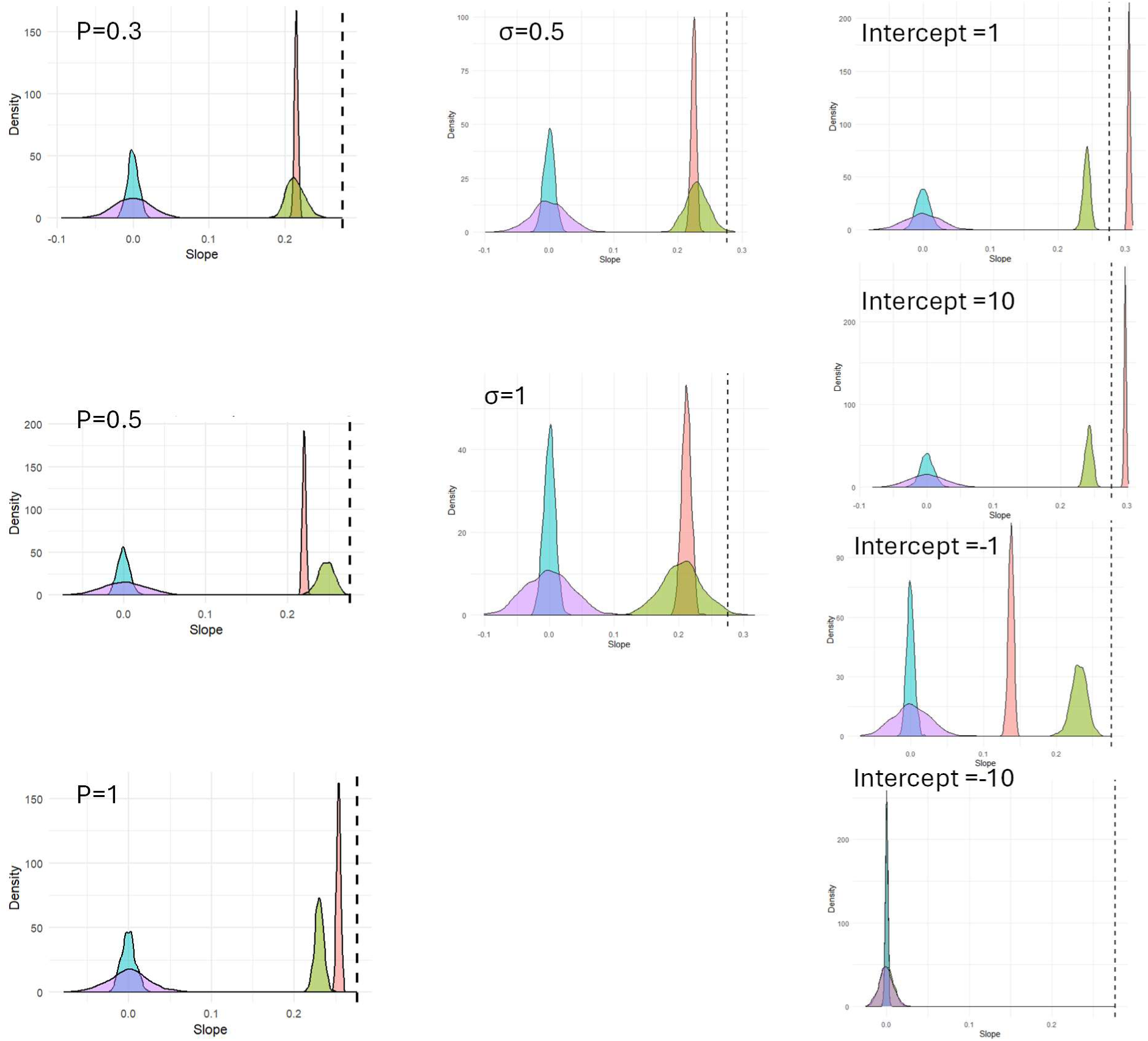
Density plots of sampled slopes across scenarios with varying prevalence, residual error (σ), and intercept, based on 1,000 simulations. In all panels, the true slope was set to 0.276. The black vertical line indicates this reference (target) slope.

**Table S1.**
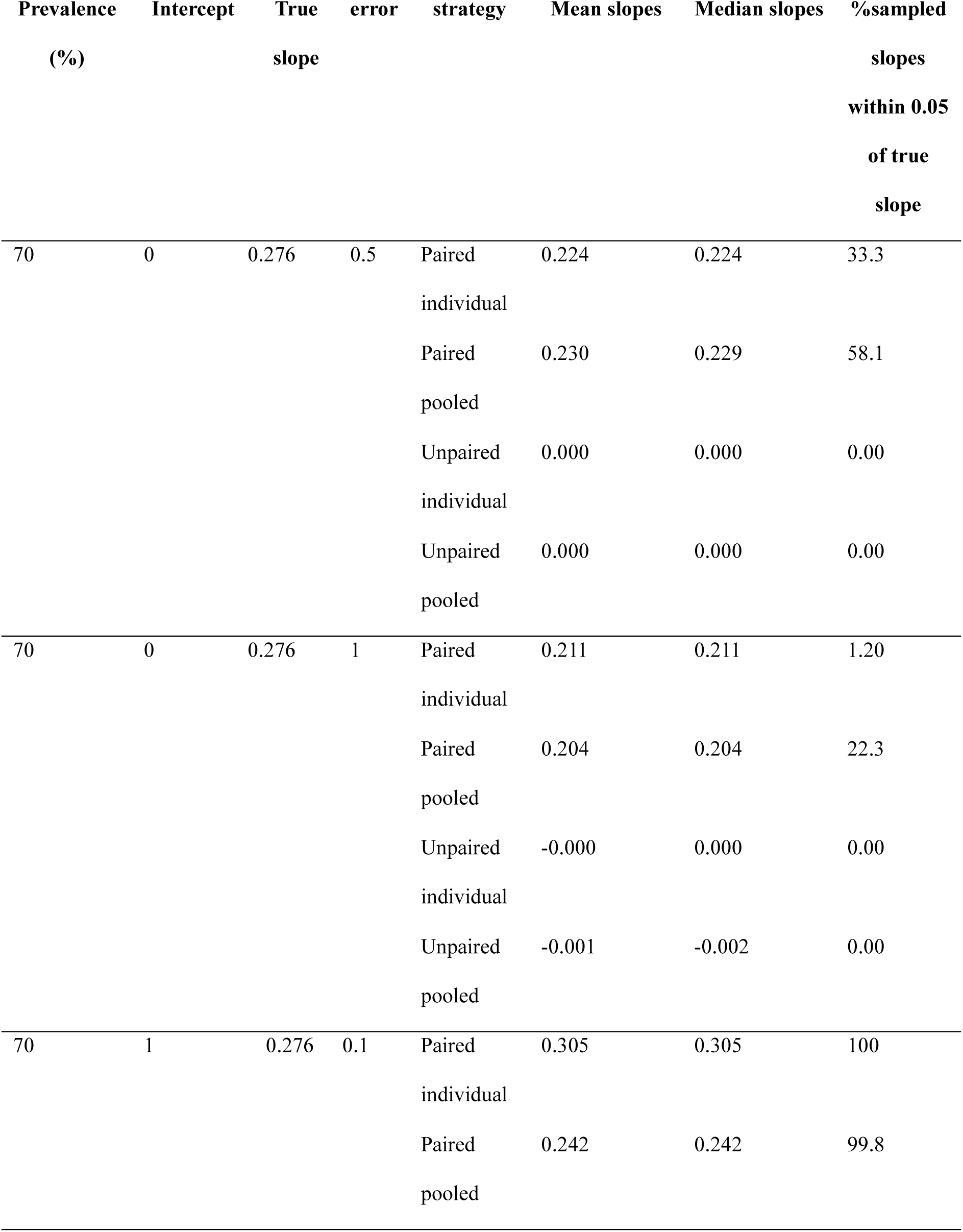
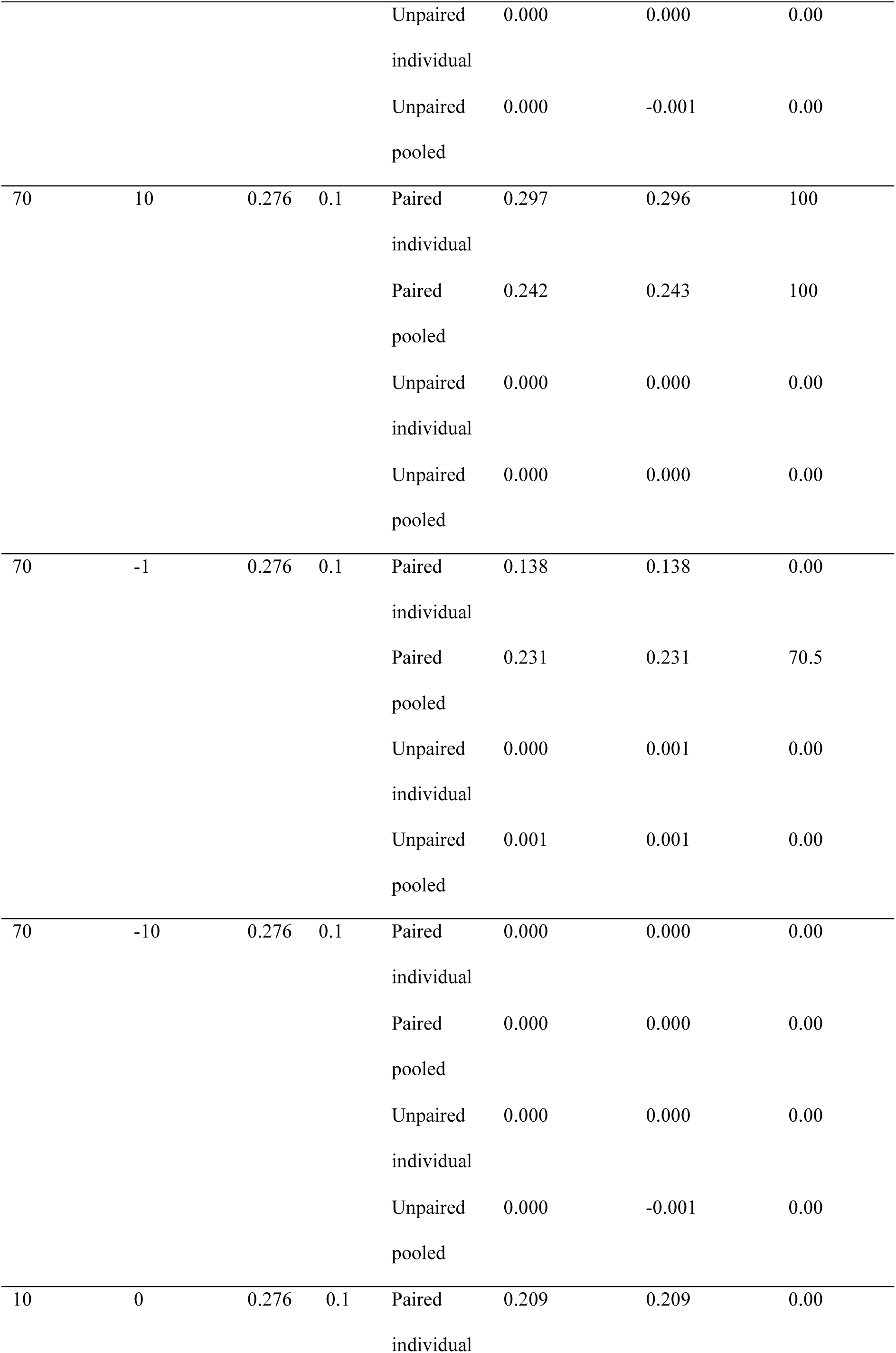
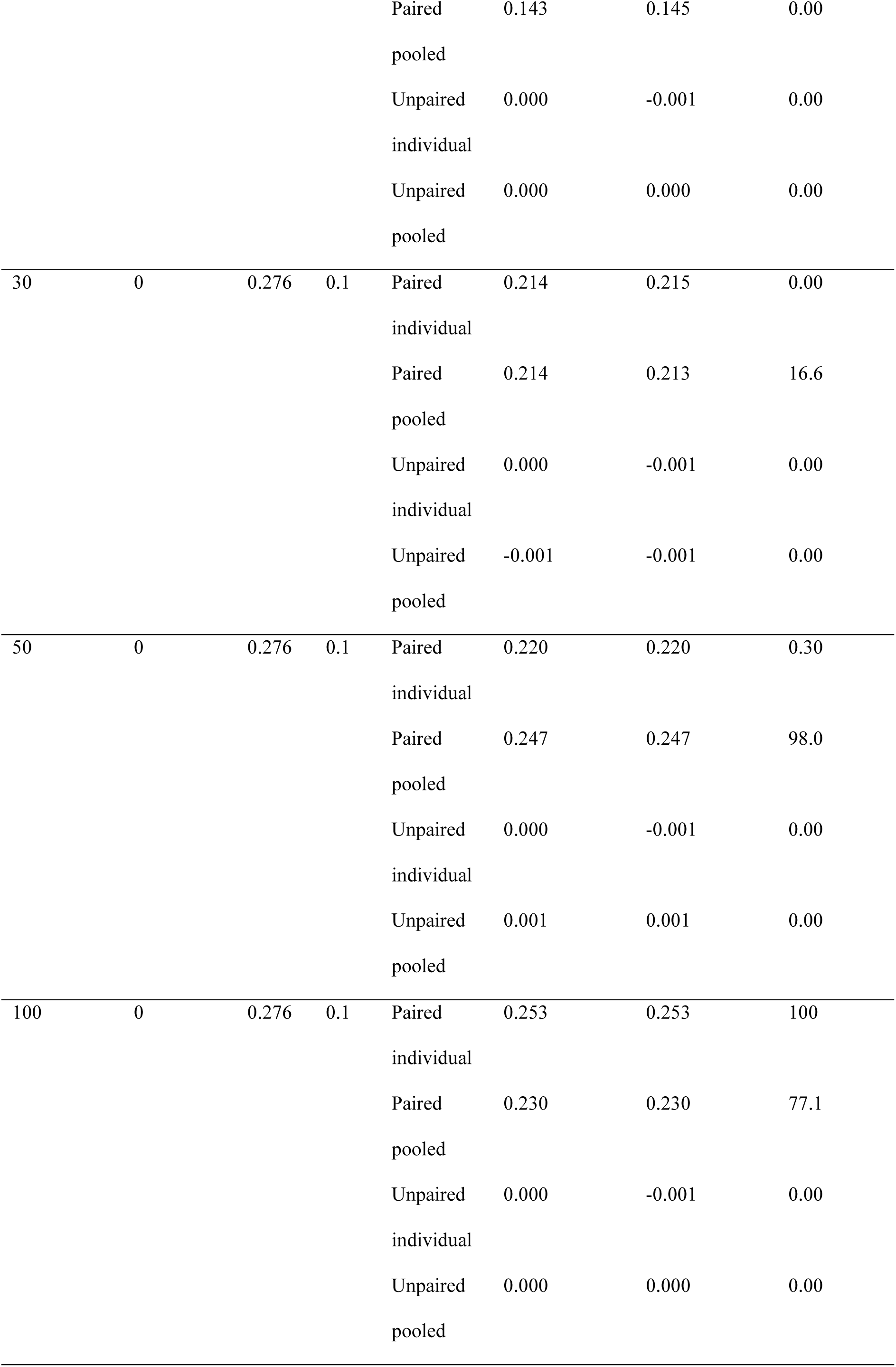
Results of scenario analysis with varying prevalence of population, intercept, and error across 100 simulations.

